# Carbon monoxide-driven proton respiration enables facultative anaerobes to survive electron acceptor limitation

**DOI:** 10.1101/2025.11.06.687094

**Authors:** Yuka Adachi Katayama, Masao Inoue, Shunsuke Okamoto, Yoshihiko Sako, Ryoma Kamikawa, Chris Greening, Takashi Yoshida

**Affiliations:** Graduate School of Agriculture, Kyoto University, Kitashirakawa Oiwake-cho, Sakyo-ku, Kyoto 606-8502, Japan; Department of Microbiology, Biomedicine Discovery Institute, Monash University, Clayton, VIC 3800, Australia; R-GIRO, Ritsumeikan University, Kusatsu, Shiga, Japan; College of Life Sciences, Ritsumeikan University, Kusatsu, Shiga, Japan

**Keywords:** carbon monoxide dehydrogenase, hydrogenase, respiration, bioenergetics, water-gas shift reaction

## Abstract

Diverse microorganisms couple the oxidation of carbon monoxide gas (CO) to the reduction of protons, producing hydrogen gas (H_2_). This energy-conserving process is mediated by the nickel-containing CO dehydrogenase/energy-converting hydrogenase (Ni-CODH/ECH). Yielding only a small supply of energy, the physiological role of this process at environmentally relevant CO levels remains unresolved. Here, we show that Ni-CODH/ECH enables metabolically flexible facultative anaerobes within the Anoxybacillaceae to survive electron acceptor limitation. Analysis of 387 Anoxybacillaceae genomes revealed that Ni-CODH/ECH had a patchy distribution and, with one exception, was mutually exclusive with the aerobic molybdenum-containing CODH. Culture experiments using the three isolates (*Parageobacillus* sp. G301, *P. thermoglucosidasius* NBRC 107763, and *Thermolongibacillus altinsuensis* B1-1) demonstrated that CO-dependent proton respiration is activated during stationary phase when exogenous electron acceptors are limiting, enhancing cell density 1.2-to 1.5-fold under 25% CO, whereas no effect was observed in a Ni-CODH knockout (Δ*cooCSF*) strain. RNA-seq analysis of strain G301 under twelve conditions revealed that Ni-CODH genes reached 2,000–12,500 TPM (top 0.2–1.9% of all genes) during stationary phase, independent of CO presence, under the predicted control of the redox-dependent transcriptional repressor Rex. Δ*cooCSF* cultures accumulated more trace CO than the wild-type, suggesting trace CO uptake by the wild-type. Thus, Ni-CODH/ECH is a redox-regulated auxiliary energy-conservation system that supports adaptation to electron acceptor limitation. Given the continual environmental supply of the two substrates for this enzyme, we propose CO-dependent proton respiration is a dependable way for metabolically flexible microorganisms to stay energized in spatiotemporally variable environments.

**Significance statement:** Microorganisms are frequently challenged to survive in environments where both energy sources and electron acceptors are limited. We reveal that facultative anaerobes activate a respiratory process that utilizes carbon monoxide (CO) and protons to survive under electron acceptor limitation. This hydrogenogenic CO-oxidizing reaction, termed CO-dependent proton respiration, is catalyzed by an ancient and minimal complex, the nickel-containing CO dehydrogenase/energy-converting hydrogenase. Although the reaction yields barely enough energy to translocate protons, this complex provides a reliable lifeline by extracting energy from trace CO and protons that are ubiquitous in nature, even when other electron acceptors are depleted. The widespread distribution of this complex across bacteria and archaea suggests that it provides a universal adaptive advantage for life under energy-limited conditions.

## Introduction

Carbon monoxide (CO) is a ubiquitous trace gas in terrestrial, marine, and host-associated environments produced through various atmospheric, biogenic, and geogenic processes (1). This gas can be harnessed by diverse bacteria and archaea as an energy and/or carbon source (1, 2). Because of its low standard redox potential (E^0′^ = –520 mV), CO oxidation can be coupled to a wide range of electron acceptors with varying redox potentials, including protons, sulfate, nitrate, and oxygen, enabling flexible energy conservation under anaerobic and aerobic conditions (3, 4). Carbon monoxide dehydrogenases (CODH) enable microorganisms to interconvert CO and CO_2_. They fall into two structurally and phylogenetically unrelated forms: nickel-containing CODH (Ni-CODH) and molybdenum-containing CODH (Mo-CODH) (4). These two enzyme types differ not only in their cofactor usage, but also in their physiological roles and redox partnerships (4). Ni-CODH is predominantly found in anaerobic microbes, where it adopts a range of roles, including CO-dependent anaerobic respiration coupled to the reduction of electron acceptors such as protons, sulfate, or nitrate, as well as mediating CO formation or direct CO uptake during the central step of acetogenesis (3–5). Predominantly found in aerobic microbes, Mo-CODH is a respiratory enzyme that supports coupling of CO oxidation to the reduction of higher potential acceptors, such as oxygen and nitrate (6). Recent studies have revealed that Mo-CODH primarily enables aerobic heterotrophs to scavenge atmospheric or trace CO, thereby sustaining cellular maintenance during carbon starvation, though some microbes can also grow autotrophically using this enzyme (1, 7). In contrast, the physiological role of CO-dependent respiration using Ni-CODH remains less defined, especially in niches with trace levels of CO.

One way that anaerobes can conserve energy is by coupling CO oxidation to proton reduction, resulting in the production of CO_2_ and hydrogen gas (CO + H_2_O ⇌ H_2_ + CO_2_; *i*.*e*. CO-dependent proton respiration, hydrogenogenic CO oxidation, or water-gas shift reaction). This is achieved through an enzyme complex formed by the association of a Ni-CODH and an energy-converting group 4 [NiFe] hydrogenase (ECH; energy-converting hydrogenase) (4), containing transmembrane subunits capable of proton pumping (8), thereby generating a transmembrane proton gradient that fuels ATP synthesis (9). This system is related to structurally characterized complexes, namely the formate hydrogenlyase (FHL) of Enterobacteriales (10) and to a lesser extent membrane-bound hydrogenases (MBH) of Thermococcales (8), that respectively couple reoxidation of endogenous formate and ferredoxin to proton reduction. It is also evolutionarily related to Complex I (8). Given the similar redox potentials of CO and H_2_, the free energy yield of CO-dependent proton respiration is one of the lowest in biology (ΔG^0′^ = –20 kJ/mol), allowing the production of less than 1 mol of ATP per reaction under standard conditions (11). Yet, hydrogenogenic CO-oxidizing activity has been reported in diverse thermophilic anaerobes and facultative anaerobes from 5 phyla, 20 genera, and 32 species as of 2020 (4). Many of these isolates harbor CO-responsive transcriptional regulators, such as *cooA* (12), *rcoM* (13), or *corQR* (14), which modulate Ni-CODH/ECH expression in response to environmental CO (4). Physiologically, under high CO concentrations, CO-dependent proton respiration has been proposed to support the chemolithoautotrophic growth of carboxydotrophs (4, 15). In addition, CO toxicity and the fast reaction kinetics of Ni-CODH (16) led to the prevailing assumption that this enzyme system acts as a CO detoxification module in CO-sensitive microbes, such as some sulfate-reducing bacteria (2, 17). However, these hypotheses do not adequately explain the physiological role of Ni-CODH/ECH in most microorganisms for four reasons: (i) many microorganisms harboring the Ni-CODH/ECH complex are obligate heterotrophs and all isolates reported to date are capable of chemoorganotrophic growth (4); (ii) many of them are found in environments where CO exists only at trace levels (e.g. soils); (iii) CODH is dispensable for CO resistance in various CO-tolerant microorganisms (7, 18, 19); and (iv) recent studies show continual aerobic respiration using atmospheric trace gases such as CO enhances survival of diverse aerobic bacteria (7, 20, 21). On this basis, we hypothesized that CO-dependent anaerobic respiration may also have a role in enhancing survival rather than detoxification.

Ni-CODH/ECH complexes was recently identified and extensively studied in *Parageobacillus thermoglucosidasius* from Anoxybacillaceae, a family of thermophilic, spore-forming, facultatively anaerobic bacteria within the phylum Bacillota, isolated from a variety of terrestrial and aquatic environments (22, 23). Notably, three species of this family (*P. thermoglucosidasius* NBRC 107763^T^, *Parageobacillus* sp. G301 and *Thermolongibacillus altinsuensis* B1-1) have been shown to mediate hydrogenogenic CO oxidation (18, 23–25). These metabolically versatile heterotrophic bacteria encode a range of respiratory machinery, including Ni-CODH/ECH, multiple [NiFe] hydrogenases, nitrate reductase, and terminal oxidases for aerobic respiration (25). CO-dependent proton respiration in the type species *P. thermoglucosidasius*, a model thermophilic fermenter for industrial application (26), has been genetically confirmed to proceed via Ni-CODH/ECH (18). Interestingly, *Parageobacillus* sp. G301 harbors both active Ni-CODH/ECH and Mo-CODH, and CO oxidation in this strain appears to be flexibly coupled to protons, nitrate, or oxygen, depending on the available electron acceptors (25, 27). However, although some strains have been found in relatively CO-rich volcanic settings (28), many members of Anoxybacillaceae are found in local soils, composts, manure, and sea sediments, where only trace levels of CO exist (22, 29). Under such conditions, the energetic contribution of the low-energy process of CO-dependent proton respiration is unclear, especially given they, like many other hydrogenogenic CO oxidizers, are capable of using a wide range of other energy sources and electron acceptors (25, 27). In addition, their genomes lack known CO-responsive transcriptional regulators (30), leaving the mechanisms of Ni-CODH/ECH gene expression and regulation largely unresolved.

To address this knowledge gap, we investigated the physiological roles and expression patterns of Ni-CODH/ECH in three Anoxybacillaceae isolates. Using a combination of genomics, culture-based physiological assays, and transcriptomic profiling under 12 conditions, we examined the timing of Ni-CODH/ECH activation in the absence of canonical CO-sensing systems. Our findings revealed that Ni-CODH/ECH-mediated hydrogenogenic CO oxidation is selectively activated during the stationary phase in the absence of external electron acceptors, potentially supplying energy for maintenance under energy-limited conditions. These results reveal a new survival strategy employed by facultative anaerobic thermophiles and suggest that Ni-CODH/ECH serves an ecological function comparable to Mo-CODH, though under anaerobic rather than aerobic conditions.

## Results

### Distribution of Ni-CODH/ECH and Mo-CODH in the family Anoxybacillaceae

To gain insight into the distribution of Ni-CODH/ECH genes, 306,260 species-level representative genomes in GlobDB release 226 (31) were initially screened (Fig. S1; Table S1). 179 genomes across 15 bacterial and 3 archaeal phyla, 55 orders, 107 genera, and 179 species encoded putative Ni-CODH/ECH genes (Fig. S1). Most of these genomes belonged to the phyla Pseudomonadota (42 genomes), Bacillota (39 genomes), Desulfobacterota (35 genomes), Methanobacteriota_B (13 genomes), and Halobacteriota (10 genomes). Putative Ni-CODH/ECH genes are encoded by genera known to use a wide range of electron acceptors, spanning microaerophiles (e.g., *Campylobacter*), methanogens (e.g., *Methanoregula*), acetogens (e.g., *Moorella*), sulfate reducers (e.g., *Desulfosporosinus*), sulfur reducers (e.g., *Thermococcus*), and iron reducers (e.g., *Geobacter*), in addition to numerous uncultivated lineages. These observations suggest that Ni-CODH/ECH complexes are not essential but may confer adaptive advantages across diverse environmental and metabolic contexts.

To better understand the roles of these enzymes, we focused on Anoxybacillaceae and analyzed all 387 available genomes (from 10 genera, 46 species) from RefSeq (Fig. 1A; Table S2). The Ni-CODH/ECH genes were consistently found to be a combination of clade F Ni-CODH and group 4a [NiFe] hydrogenase (Hyc/Hyf type). They were detected in 28 genomes, all from the genera *Parageobacillus* and *Thermolongibacillus*, including all 26 genomes of the species *P. thermoglucosidasius* (except for two genomes annotated as *P. thermoglucosidasius*_A) and one of the two genomes of *T. altinsuensis*. In addition, 19 Anoxybacillaceae within the genera *Parageobacillus* and *Saccharococcus* encode the aerobic Mo-CODH (encoded by *coxLMS*). Among them, 15 of the 21

**Fig. 1.**
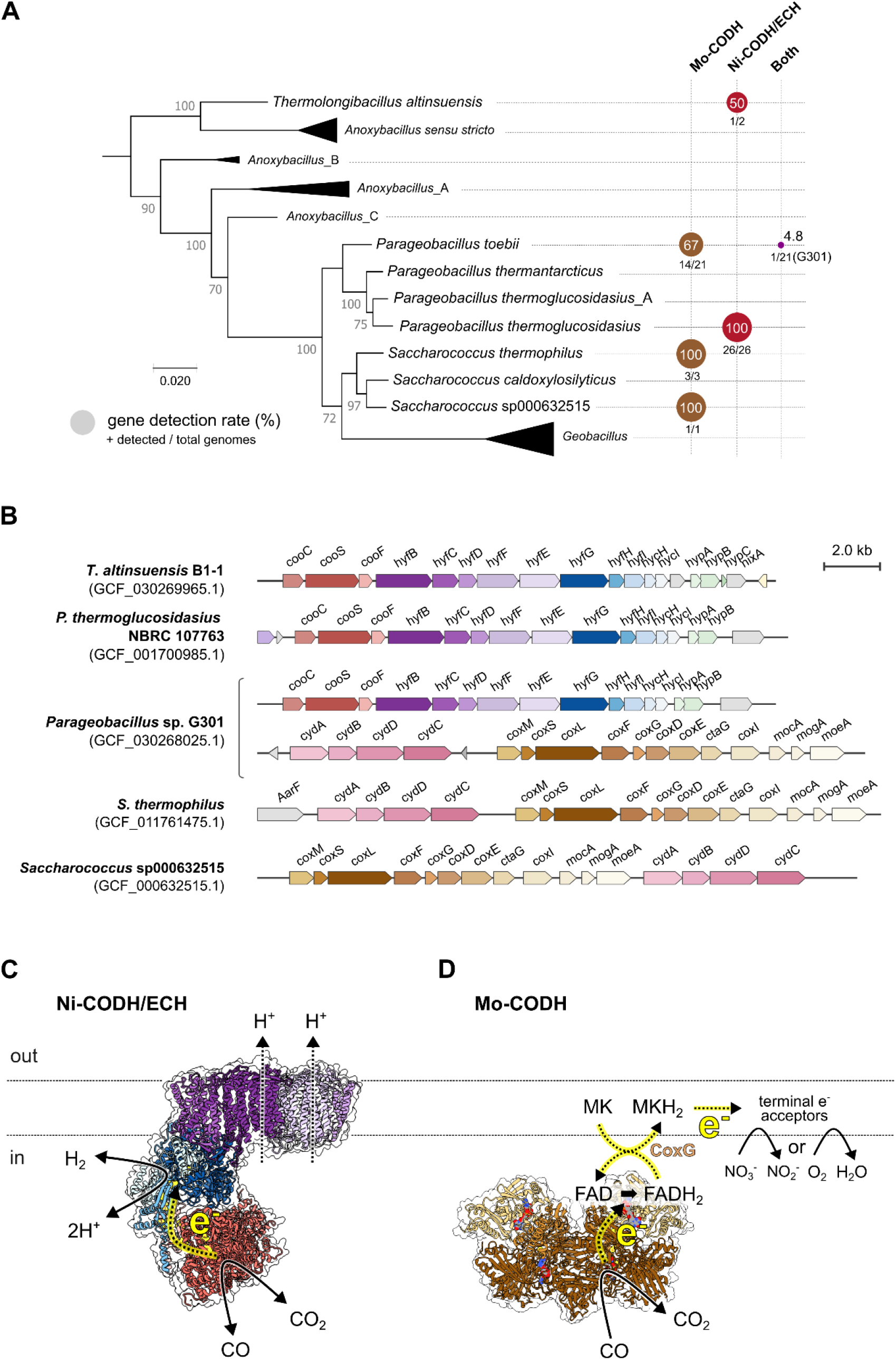
Distribution, genetic organization, and structural models of CODHs in the family Anoxybacillaceae. (A) A maximum-likelihood phylogenetic tree was constructed based on 120 concatenated single-copy core genes using the GTDB backbone tree (bac120_r226). The tree includes representative genomes from the Anoxybacillaceae family, with major clades collapsed and labeled based on the GTDB classification. Bootstrap support values (out of 100) are shown at key nodes. Scale bar represents 0.02 substitutions per site. On the right, gene detection rates of CODH genes are shown for each genus or species. Circles indicate the proportion of genomes within each clade encoding (i) Mo-CODH (brown), (ii) Ni-CODH/ECH (red), or (iii) both (purple). Gene detection rates were expressed as percentages, with circle sizes scaled accordingly. Detection rates were calculated from all available RefSeq genomes of the Anoxybacillaceae family. The numbers below the circles indicate the total number of genomes analyzed and the number of genomes in which each gene was detected. (B) Genetic organization of representative genomes encoding Ni-CODH/ECH and/or Mo-CODH. Gene abbreviations: *coo* (Ni-CODH subunits and a maturation factor), *hyf* (ECH hydrogenase and transmembrane antiporter-like subunits), *cox* (Mo-CODH subunits and maturation factors), and *cyd* (cytochrome ubiquinol oxidase modules). Predicted protein structure and schematic model of (C) Ni-CODH/ECH and (D) Mo-CODH from *Parageobacillus* sp. G301, generated using AlphaFold 3. Dotted lines with yellow highlights in the structures indicate the predicted electron flows.

*P. toebii* genomes, all three *Saccharococcus thermophilus* genomes, and one genome of *Saccharococcus* sp000632515 contain *coxL*. Surprisingly, the two types of CODH genes exhibited a patchy and largely mutually exclusive distribution across the family regardless of phylogenetic relatedness (Fig. 1A). Only *Parageobacillus* sp. G301, classified within *P. toebii*, encodes genes for both the Ni-CODH/ECH and Mo-CODH, as reported previously (25). The putative operons encoding both complexes have near-identical organizations across the Anoxybacillaceae (Fig. 1B). The list of analyzed genomes, their GTDB taxonomy, and the distributions of *cooS*/*hyfG* (catalytic subunit genes for Ni-CODH/ECH) and *coxL* are provided in Table S2.

Structural modelling suggests Ni-CODH/ECH and Mo-CODH generate proton-motive force in *Parageobacillus* sp. G301 through distinct mechanisms (Fig. 1C, D). The Ni-CODH/ECH complex is predicted to form a minimalistic respiratory chain formed by four interacting modules: (i) a peripheral Ni-CODH module that inputs CO-derived electrons (CooS), (ii) a ferredoxin-like protein that transfer electrons between the two catalytic enzymes (CooF), (iii) a [NiFe]-hydrogenase module that uses these electrons to reduce protons to H_2_ (HyfGHI), and (iv) a transmembrane domain that uses the energy released by electron transfer to pump protons across the membrane (HyfBCDFE). The second, third, and forth modules are closely related to other energy-converting complexes, especially FHL (10). In contrast, the Mo-CODH enzyme (CoxLMS) adopts a near identical structure to those described in other aerobic bacteria (32) and forms part of multi-component respiratory chain, in which electrons are first transferred to quinones via CoxG and then to terminal oxidases or nitrate reductases.

### Stationary-phase hydrogenogenic CO oxidation in three Anoxybacillaceae isolates

Because Ni-CODH/ECH and Mo-CODH are largely distributed in a mutually exclusive manner, we considered whether these two CODH systems might have comparable physiological functions in CO utilization. Mo-CODH enables aerobic heterotrophs to scavenge trace amounts of CO, supporting their survival over extended periods when nutrients are scarce (1, 7). To examine this possibility, we investigated the CO oxidation and growth patterns in three thermophilic facultative anaerobes: *Parageobacillus* sp. G301 (encoding both Mo-CODH and Ni-CODH/ECH) (25), *P. thermoglucosidasius* NBRC 107763^T^ wild-type (WT; Ni-CODH/ECH-positive) (29) and its Ni-CODH markerless gene disruptant (Δ*cooCSF*) (18), and *T. altinsuensis* B1-1 (Ni-CODH/ECH-positive) (24).

All three strains exhibited comparable growth under CO-free anaerobic conditions, reaching mean OD_600_ values of 0.028–0.049 in minimal medium containing sodium pyruvate as the sole organic carbon source (Figs. 2A–C). In the presence of 25% CO + 75% N_2_, growth proceeded similarly during the exponential phase, but after entering the stationary phase (10 h), all strains showed an increase in cell density, reaching OD_600_ values 1.5, 1.5, and 1.2-fold higher than those in CO-free conditions by 70 h in the stains G301, B1-1, and *P. thermoglucosidasius* WT, respectively. For *Parageobacillus* sp. G301, OD_600_ initially decreased after 9 h (from 0.028 to 0.016), but subsequently increased significantly to 0.041 only in the presence of CO (*p* < 0.001), indicating CO-dependent secondary growth (Fig. 2A). *P. thermoglucosidasius* WT also exhibited a slow but continuous increase under CO (from 0.031 at 9 h to 0.071 at 70 h; *p* < 0.05) (Fig. 2B). This CO-dependent stationary-phase growth enhancement was absent in the Δ*cooCSF* mutant of *P. thermoglucosidasius* (*p* < 0.05; Fig. 2B), indicating an Ni-CODH-dependent process, consistent with a previous study (18). *T. altinsuensis* B1-1 slightly decreased in OD_600_ under CO-free conditions (from 0.029 to 0.018), but maintained higher cell density when CO was supplied (from 0.034 to 0.029), suggesting a weak CO-stimulated growth (Fig. 2C). Gas measurements revealed that CO oxidation was negligible during exponential growth and instead initiated during stationary phase (Figs. 2D–F). The amount of CO steadily declined from 2.0–2.1 mmol to below detection limit after 21–40 h, concomitant with H_2_ accumulation and CO_2_ production to 2.2–2.4 mmol and 1.6–2.1 mmol, respectively (Figs. 2D–F). This indicates Ni-CODH/ECH-mediated hydrogenogenic CO oxidation at the expected equimolar stoichiometry.

**Fig. 2.**
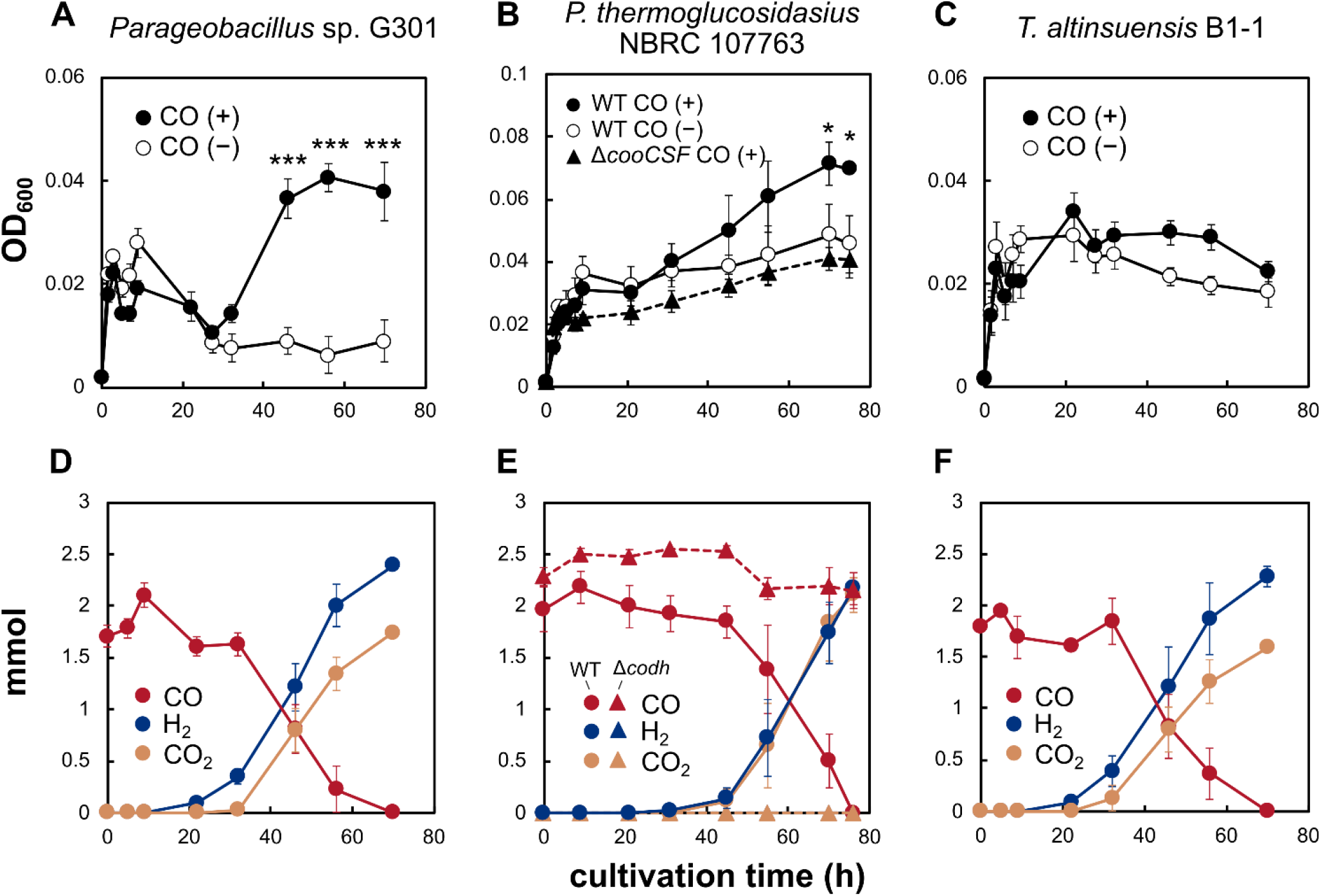
Growth of the three thermophilic facultative anaerobes from Anoxybacillaceae family in the presence and absence of CO. OD_600_ (black lines) of *Parageobacillus* sp. G301 (A), *P. thermoglucosidasius* NBRC 107763 wild-type and Δ*cooCSF* (B), and *Thermolongibacillus altinsuensis* B1-1 (C) was monitored under either 25% CO + 75% N_2_ (closed symbols) or 100% N_2_ (open symbols) in a minimal medium supplemented with 15 mM sodium pyruvate. The gas composition of the headspace of the bottles was measured in the culture of *Parageobacillus* sp. G301 (D), *P. thermoglucosidasius* wild-type and Δ*cooCSF* (E), and *T. altinsuensis* B1-1 (F) containing 25% CO + 75% N_2_. The figures show the total amounts of CO (red), CO_2_ (yellow), H_2_ (blue). The experiment was performed in triplicate. The error bars represent the standard error of the mean. Statistical analysis was performed using a linear mixed-effects model for repeated measures (\*\*\**p* < 0.001, \**p* < 0.05).

*Parageobacillus* sp. G301 is currently the only identified member of the Anoxybacillaceae family that encodes both active Ni-CODH/ECH and Mo-CODH (25). In this strain, Mo-CODH-dependent CO oxidation is thought to be coupled with nitrate reduction and aerobic respiration (25, 27). To examine the role of Mo-CODH, strain G301 was cultured in a minimal medium with 5 mM sodium pyruvate (Fig. 3; Fig. S2). Growth rates during the exponential phase were similar between cultures with and without CO under both nitrate-supplemented (25 mM potassium nitrate, 100% N_2_) and oxygen-supplemented (25% CO + 15% O_2_ + 60% N_2_) conditions. In nitrate-supplemented cultures, CO led to an increase in OD_600_ from 0.046 at 5 h to 0.103 at 8 h, whereas CO-free cultures reached only 0.078 (Fig. 3A). This 1.3-fold higher cell density was accompanied by continuous CO oxidation (Fig. 3B). No H_2_ was detected under nitrate-respiring conditions, indicating that nitrate suppresses CO-dependent proton respiration, as observed in *P. thermoglucosidasius* (27). These results suggest that, in addition to supporting aerobic survival (7), Mo-CODH can support anaerobic survival through coupling to nitrate reduction. In contrast, under O_2_-respiring conditions, OD_600_ in CO-amended cultures became higher than in CO-free controls during the late-log to early-stationary phase (0.069 vs. 0.051 at 8 h) but became nearly identical when CO oxidation temporarily ceased (22–31 h) (Fig. S2A). Subsequently, when CO oxidation resumed (31–46 h), during which 0.27 mmol of CO was oxidized and 0.16 mmol of H_2_ was produced, OD_600_ remained stable around 0.07 in CO-amended cultures, whereas it declined from 0.087 to 0.048 in CO-free cultures (Fig. S2B). These observations indicated that G301 initially utilized CO through aerobic oxidation using Mo-CODH, followed by hydrogenogenic CO oxidation using Ni-CODH/ECH once O_2_ was depleted, supporting cell maintenance during the late stationary phase.

**Fig. 3.**
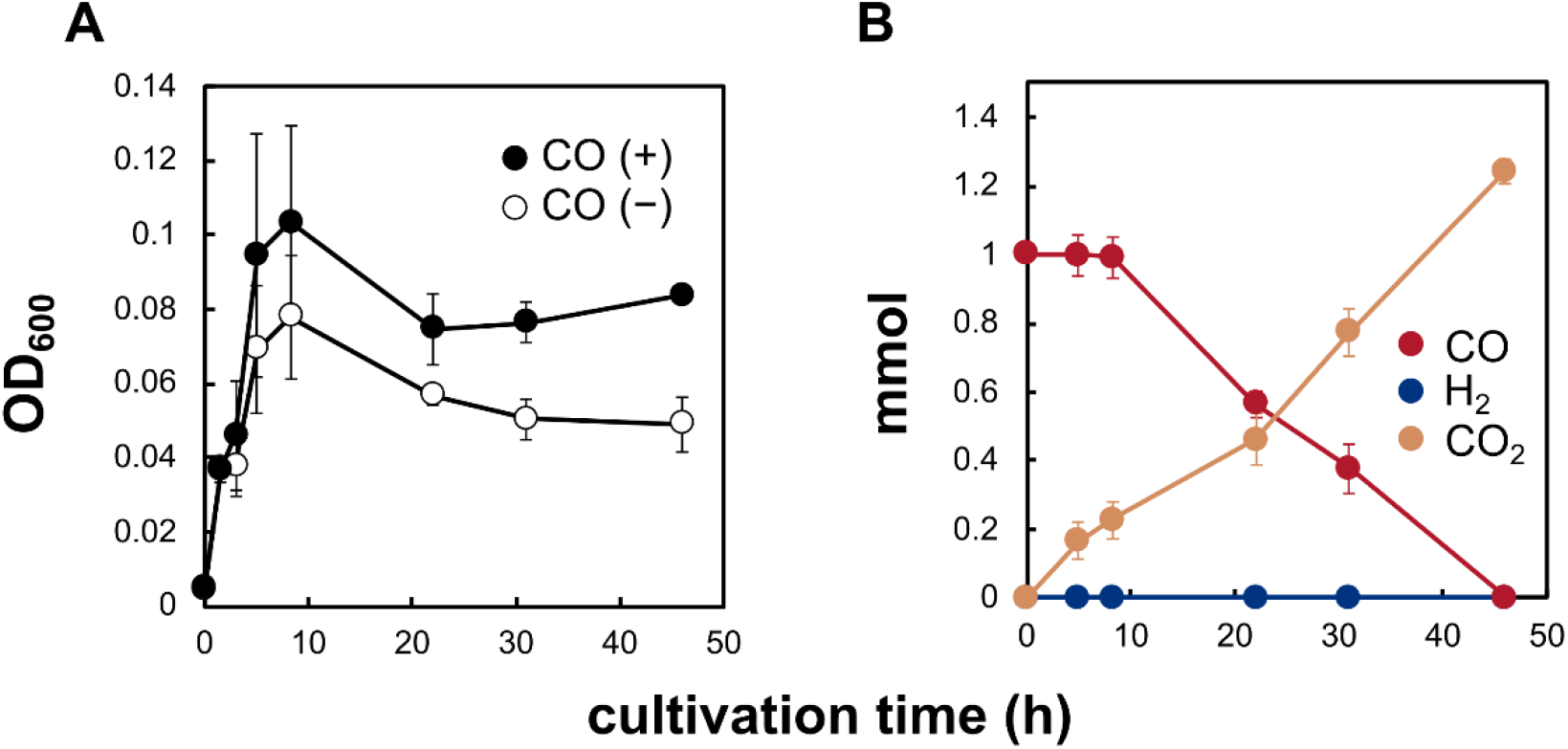
Growth and gas composition of *Parageobacillus* sp. G301 under nitrate-respiring conditions with or without CO. Cultures were grown in basal medium supplemented with 5 mM sodium pyruvate under either 25 mM KNO_3_ in the presence of CO (closed symbols) or in its absence (open symbols). (A) Growth curves (OD_600_). (B) Total amounts of CO (red), CO_2_ (yellow), H_2_ (blue) during the same incubation. The error bars represent the standard error of the mean.

### Ni-CODH/ECH expression is CO-independent and activated under fermentative conditions

Although the culture experiments revealed that CO oxidation mediated by both Ni-CODH/ECH and Mo-CODH occurs and enhances cell density during the late-log to stationary phase, the regulatory patterns of these enzymes remain unclear. To compare the transcriptional patterns of Ni-CODH/ECH and Mo-CODH, we performed RNA-seq on *Parageobacillus* sp. G301, the only known isolate encoding both Ni-CODH/ECH and Mo-CODH (25) under 12 different conditions (Fig. 4). Cells were cultured in minimal medium supplemented with 0.1% yeast extract (potential carbon and energy sources), either 25 mM sodium pyruvate (carbon and energy sources) or 20% CO (energy source), and three electron acceptors (oxygen, nitrate, or no exogenous acceptors, *i*.*e*. protons). Log-phase samples were collected when the OD_600_ reached 0.1, whereas stationary-phase samples were taken at the first time point when the OD_600_ no longer increased. The growth and gas profiles under each condition are shown in Fig. 4A–F. Consistent with previous physiological experiments (Fig. 2, 3; Fig. S2) (25, 27), hydrogenogenic CO oxidation was evident only under electron acceptor-free conditions (Fig. 4F). Principal component analysis (PCA) revealed a clear separation of transcriptomic profiles according to both electron acceptor type and growth phase, with oxygen- and nitrate-respiring conditions clustering distinctly from exogenous electron acceptor-free conditions (Fig. 4G).

**Fig. 4.**
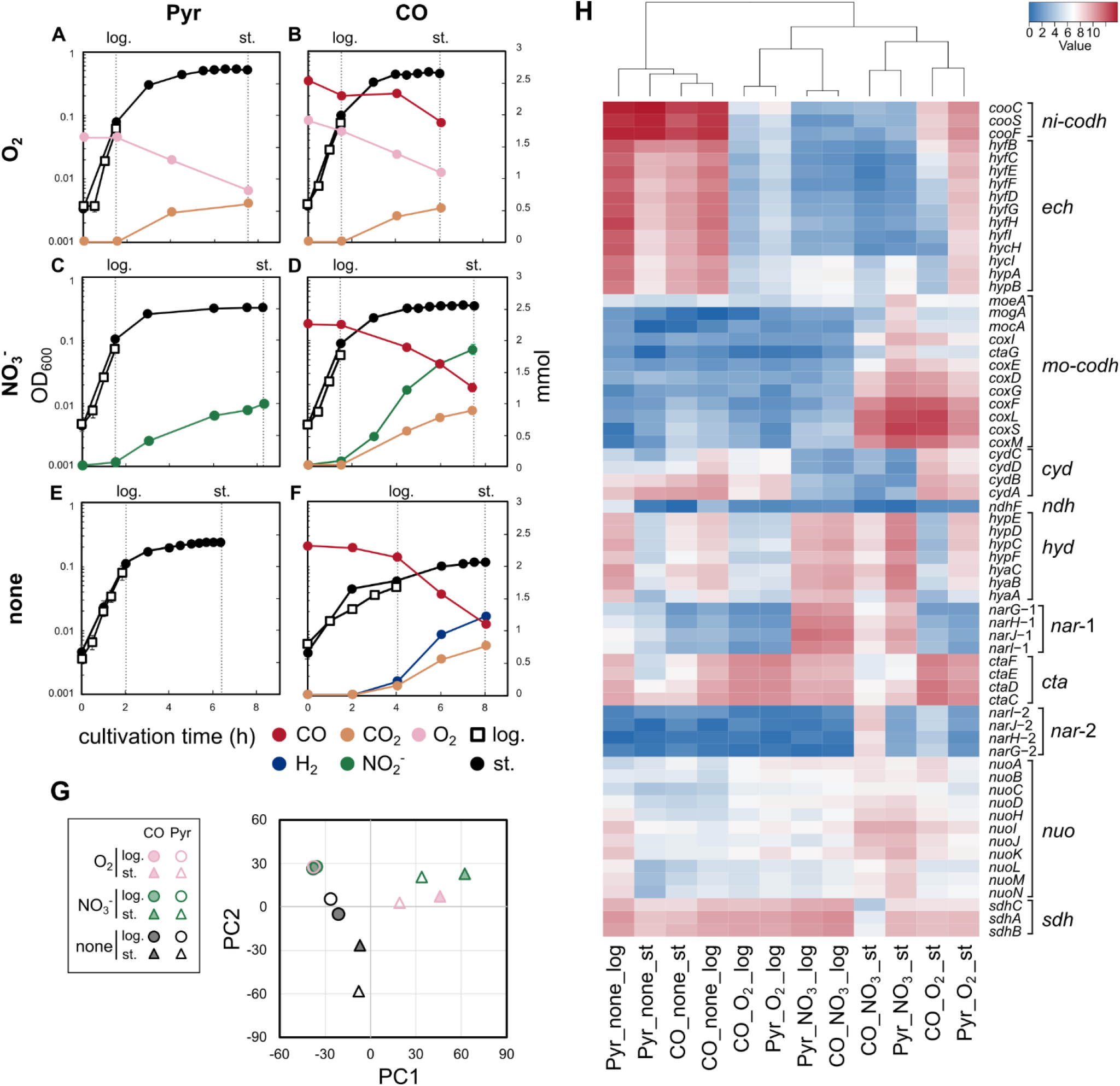
Transcriptomic analysis of *Parageobacillus* sp. G301 under twelve different carbon and electron acceptor conditions. (A–F) Growth and headspace gas composition of the strain G301 cultured under various electron acceptor conditions with either 25 mM pyruvate under 100% N_2_ (Pyr, A, C, E) or 20% CO + 80% N_2_ (CO, B, D, F) in the minimal media containing 0.1% yeast extract. OD_600_ (black circles or open squares), total gas amounts (colored circles), and cultivation time are shown. Electron acceptor conditions: (A–B) oxygen (O_2_, pink), (C–D) nitrate (NO_3_−, green), (E–F) no external electron acceptor. Open squares indicate time points of RNA sampling at logarithmic (log.) phases, and closed circles those at stationary phase (st.). The error bars represent the standard error of the mean. (G) Principal component analysis (PCA) of transcriptomic profiles across all conditions and time points. Conditions and phases are distinguished by color and shape, respectively. (H) Heatmap showing the transcriptional levels, log_2_ (TPM + 1), of selected genes involved in nickel-containing carbon monoxide dehydrogenase (*ni-codh*), energy-converting hydrogenase (*ech*), molybdenum-containing carbon monoxide dehydrogenase (*mo-codh*), cytochrome ubiquinol oxidase (*cyd*), group 1d [NiFe]-hydrogenase (*hyd*), nitrate reductase (*nar*-1, *nar*-2), cytochrome *c* oxidase (*cta*), NADH dehydrogenase (*nuo, ndh*), and succinate dehydrogenase (*sdh*). Samples are clustered by hierarchical clustering (top dendrogram).

Next, we examined the relative transcript levels of the respiratory and CODH-related genes (Fig. 4H). The transcriptional patterns of Ni-CODH/ECH genes and Mo-CODH genes were markedly different. All 15 Ni-CODH/ECH-related genes (*cooCSF*, QSJ37_RS02060–2070; *hyfBCEFDGHIhycHIhypAB*, QSJ37_RS02060–2130) showed similar transcriptional patterns and were grouped to the same cluster (Table S3). Their relative transcript levels were strongly elevated under external electron acceptor-free conditions, independent of CO supplementation (Fig. 4H; Table S3). Under the four electron acceptor-free conditions, the catalytic subunit gene *cooS* (QSJ37_RS02065) reached TPM values of 2,271–12,503, ranking within the top 0.2–1.9% of all genes (Table S3). Ni-CODH/ECH genes expression also increased during the stationary phase under oxygen-supplemented conditions, with *cooS* TPM values rising from 25 and 53 at log-phase to 275 and 890 at stationary phase without and with pyruvate, respectively (|log FC| = 3.6–4.0, FDR < 0.01) (Fig. S3). These transcriptional dynamics are in agreement with previous reports showing that *P. thermoglucosidasius* initiates Ni-CODH/ECH-dependent CO oxidation immediately after aerobic respiration ceases (23, 27). In the presence of nitrate, *cooS* TPM values were consistently low (4.0– 12.7) (Fig. 4H), suggesting that CO oxidation by Ni-CODH is not coupled with nitrate reduction in *Parageobacillus* sp. G301, consistent with the repression of anaerobic CO oxidation in *P. thermoglucosidasius* in the presence of nitrate (Fig. 3, 4D) (27). G301 harbors two dissimilatory nitrate reductase complexes (NarGHJI), with two NarG genes (*narG*-1, QSJ37_RS03390 and *narG*-2, QSJ37_RS07450) coding for the catalytic alpha subunit. Among them, only *narG*-2 showed transcriptional upregulation with CO under nitrate-respiring conditions, with stationary-phase TPM values increasing from 8.5 to 240 (|log FC| = 5.5, FDR < 0.01; Fig. 3H; Fig. S3), suggesting the potential for CO responsiveness.

Genes encoding Mo-CODH (*coxLMSG*) exhibited different expression patterns from those of the Ni-CODH/ECH genes. As reported for other Mo-CODH-containing bacteria (7), Mo-CODH genes expression increased during starvation-induced stationary phase under both oxygen- and nitrate-respiring conditions, regardless of CO supplementation (Fig. 4H; Table S3). For example, *coxL* (QSJ37_RS02820) TPM values significantly increased from 6.3 and 4.2 (log-phase) to 5,736 and 1,232 (stationary phase) under oxygen-supplemented conditions and from 19 and 13 to 2,101 and 4,801 under nitrate-respiring conditions in the absence and presence of pyruvate, respectively (FDR < 0.01; Fig. S3). These patterns suggest that Mo-CODH and Ni-CODH/ECH are both expressed to enhance cellular energy supply in response to limitation of organic energy sources and respiratory electron acceptors respectively. The relative transcriptional level of cytochrome ubiquinol oxidase (*cydAB*, QSJ37_RS02850, QSJ37_RS02855) was low in the presence of nitrate, with TPM values of 4.3–22.2 (Fig. 4H).

Next, global transcriptional patterns were analyzed using hierarchical clustering (Table S3), and genes involved in central metabolism were mapped (Fig. 5). The putative operon encoding Ni-CODH/ECH genes are clustered with 19 other genes, including L-lactate dehydrogenase gene (*ldh*), L-lactate permease gene, and genes involved in branched-chain amino acid metabolism such as acetolactate synthase genes (Table S3). These genes showed significant upregulation exclusively under electron acceptor-free conditions, regardless of the carbon or electron donor source. Similar to *cooS, ldh* reached TPM values of 363–1,327 under four conditions without electron acceptors, with a mean log FC change of 4.9 compared to the other eight conditions (Table S3). During electron acceptor limitation, cells face the simultaneous challenge of disposing excess reductant and producing ATP. By co-expressing two key enzymes, *Parageobacillus* sp. G301 can simultaneously dispose of NADH by producing the fermentative end-product L-lactate by LDH, while maintaining ATP production by Ni-CODH/ECH. As depicted in the metabolic reconstruction (Fig. 5), under electron acceptor limited conditions, pyruvate-grown cells also transcribe genes involved in the oxidation of pyruvate to CO_2_ via the TCA cycle, as well as those for the fermentative production of formate, ethanol, and acetate, likely to balance electron flux and energy requirements. Acetate production is likely a primary source of ATP supplemented by inputs from the Ni-CODH/ECH complex.

**Fig. 5.**
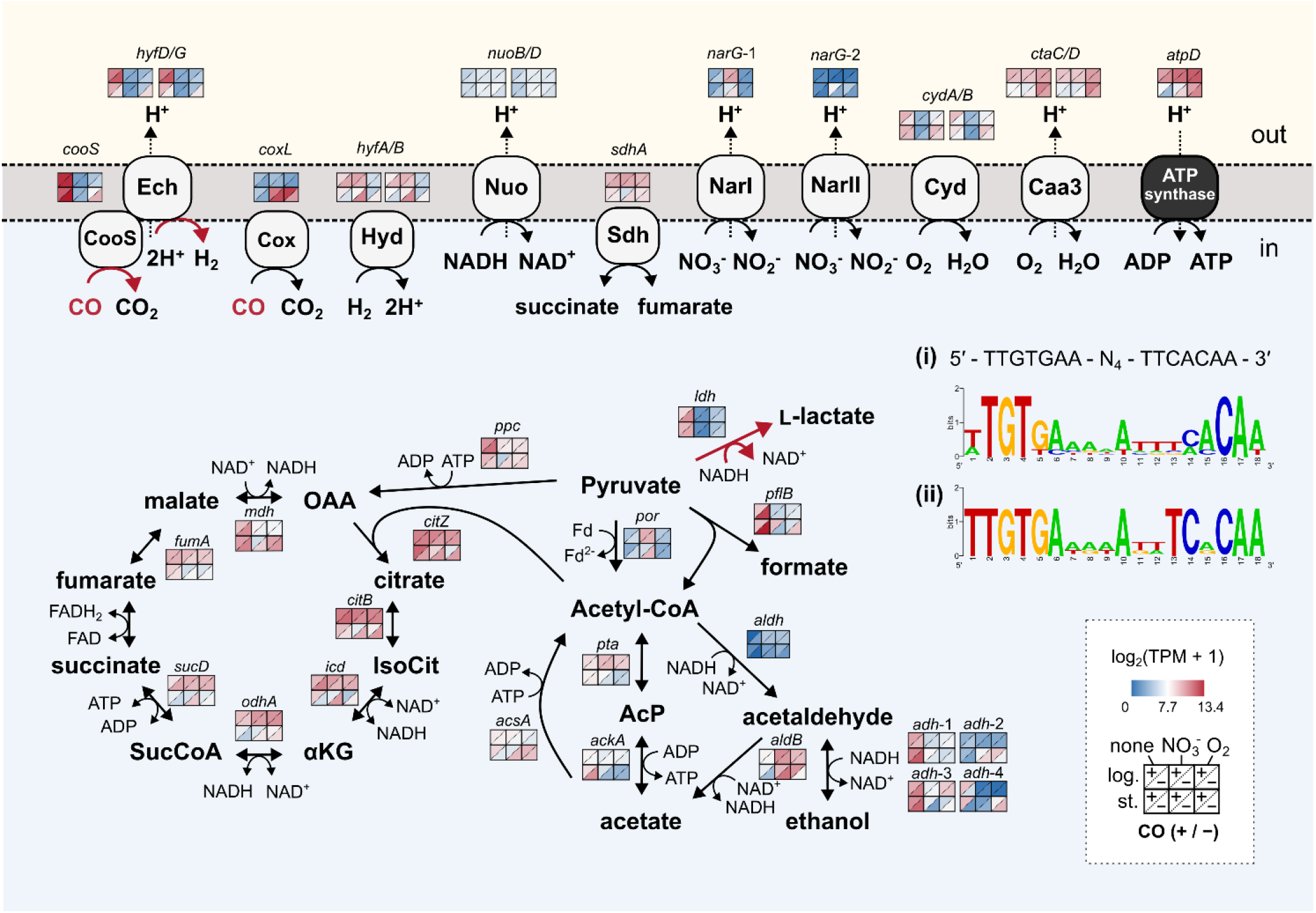
Transcriptional landscape of central metabolism and respiratory pathways under different conditions. This schematic summarizes the gene expression patterns in the key metabolic and respiratory pathways of *Parageobacillus* sp. G301 across the 12 experimental conditions, varying by exogenous electron acceptor availability (none, NO_3_^−^, O_2_), carbon source (pyruvate or CO), and growth phase (log or stationary). Expression levels are indicated for each gene by a heatmap showing log_2_ (TPM +1) values, with red representing higher expression and blue representing lower expression. Each heatmap cell indicates the following conditions: columns represent electron acceptor conditions (left: none, middle: NO_3_^−^, right: O_2_), rows indicate the growth phase (top: log-phase, bottom: stationary phase), and within each cell, the upper-left triangle corresponds to the absence of CO, and the lower-right triangle corresponds to the presence of CO. Black arrows indicate enzymatic reactions or gene products. Genes and complexes related to carbon monoxide oxidation (CooS, Cox), hydrogen metabolism (Hyd, group 1d [NiFe] hydrogenase; Ech), electron transport chain (Nuo, Nar, Sdh, Cyd, Caa3, ATP synthase), TCA cycle, and mixed acid fermentation are shown. The abbreviations represent as follows: OAA, oxaloacetate; SucCoA, succinyl-CoA; αKG, alpha ketoglutarate; IsoCit, isocitrate; AcP, acetyl-phosphate. Genes significantly upregulated under all four conditions lacking external electron acceptors are indicated by red arrows. Predicted Rex-binding motifs were also identified upstream of redox-regulated genes. Rex is a redox-sensing transcriptional repressor that binds DNA as a homodimer when the intracellular NADH/NAD^+^ ratio is low and dissociates upon NADH binding, thereby derepressing target genes. Panel (i) shows the predicted Rex-binding motif among genes previously recognized as Rex-regulated in *Staphylococcus aureus* and *Bacillus subtilis*, including *ldh, pflB*, and *cydA*. Panel (ii) illustrates the predicted Rex-binding motif upstream of Ni-CODH/ECH (*cooC*) genes across *Parageobacillus* sp. G301, *P. thermoglucosidasius*, and *T. altinsuensis* B1-1, demonstrating conserved sequence patterns consistent with the consensus motif (5′-TTGTGAA-N_4_-TTCACAA-3′). Details of the Rex-binding sites are shown in Fig. S4.

Given that *ldh* is a well-characterized target of the redox-sensing transcriptional regulator Rex (or YdiH) in *Bacillus subtilis* and related Bacillota (33–37) (Fig. S4) and that the strain G301 possesses putative Rex-encoding gene (QSJ37_RS17140), we searched for conserved regulatory elements upstream of these genes. Comparative motif analysis identified a previously reported generalized Rex-binding sequence (TTGTGAA-N_4_-TTCACAA)-like motif (36) located in the upstream of previously reported Rex-regulated genes such as *cydA, pflB* (pyruvate-formate lyase gene), *narG*, and *ywcJ* (nitrite transporter gene), as well as 32 and 36 bp upstream of both *ldh* and *cooCSF*, respectively (Fig. 5 (i); Fig. S4). Furthermore, the Rex-binding-like motif was also found 37 bp and 31 bp upstream of the putative operon encoding Ni-CODH/ECH in *P. thermoglucosidasius* and *T. altinsuensis*, respectively (Fig. 5 (ii); Fig. S4).

### Ni-CODH recycles trace levels of CO produced under anaerobic conditions

*Parageobacillaceae* are likely to scavenge CO at a range of concentrations produced through various processes. While atmospheric CO scavenging is feasible only using Mo-CODH, given CO-dependent proton respiration is thermodynamically unfavorable at atmospheric CO concentrations (as per the Nernst equation), Ni-CODH potentially scavenges elevated levels of CO produced during biotic and abiotic processes. Following recent reports on CO production by *Bacillus* and *Geobacillus* species in compost (38, 39), we tested whether *Anoxybacillaceae* strains with varying CODH genotypes could also produce CO (Fig. S5). Under anaerobic conditions, CO levels in minimal medium remained below the detection limit (0.5 ppm) (data not shown), whereas 2.5–5.4 ppm CO was detected in rich TGP medium (Fig. S5A), indicating abiotic formation from medium components. CO levels were significantly elevated in a *P. thermoglucosidasius* mutant unable to consume CO (Δ*cooCSF*), reaching 29.5 ± 6.2 ppm after 19 h. These results suggest both abiotic and potentially biotic CO generation occur under anaerobic conditions. In contrast, CO concentrations in *P. thermoglucosidasius* wild-type and *Parageobacillus* sp. G301 cultures remained similar to the control. This suggests that Ni-CODH reoxidizes the environmentally relevant nanomolar concentrations of CO produced.

## Discussion

This study identifies a previously unrecognized physiological role of anaerobic CO oxidation in thermophilic facultative anaerobes. Hydrogenogenic CO oxidation occurred during the stationary phase under external electron acceptor limitation, indicating activation under fermentative conditions. All three tested strains mediated hydrogenogenic CO oxidation when no exogenous electron acceptors were available. The magnitude of this effect varied among strains: *Parageobacillus* species exhibited a secondary increase in the optical density concurrent with hydrogenogenic CO oxidation, suggesting that the energy derived from CO is utilized for biomass synthesis, while *T. altinsuensis* B1-1 maintained cell density without additional growth under 25% CO. However, this condition far exceeds environmental CO levels where growth stimulation is expected to be minimal. Activation of CO oxidation under energy-depleted conditions parallels the physiological role of atmospheric CO and H_2_ oxidation, the former mediated by Mo-CODH (1, 6, 7, 40). Notably, Ni-CODH/ECH and Mo-CODH exhibited largely mutually exclusive distributions (Fig. 1, Fig. S1). Furthermore, strain G301 mediated three types of CO-driven respiration (proton, nitrate, and aerobic respiration) during stationary phase, mediated either by Ni-CODH or Mo-CODH (Fig. 2, 3), indicating that CO is consistently utilized as a supplementary energy source during energy deprivation under various redox potentials. Therefore, we propose that Ni-CODH/ECH performs a function analogous to that of Mo-CODH, but is specifically adapted to strictly anaerobic environments and low redox conditions. CO-dependent proton reduction via Ni-CODH/ECH yields substantially less energy (ΔG^0′^ = −20 kJ/mol) than CO oxidation coupled with respiratory electron acceptors, such as O_2_ (ΔG^0′^ = −258 kJ/mol) or nitrate (ΔG^0′^ = −181 kJ/mol). Nonetheless, in anaerobic environments where external electron acceptors are absent, Ni-CODH/ECH might provide a dependable means to generate a proton-motive force using cellular protons as the electron sink and endogenous or environmentally produced CO as an electron donor.

Transcriptional patterns further indicate that Ni-CODH/ECH expression responds to intracellular redox balance. In *Parageobacillus* sp. G301, Ni-CODH/ECH gene cluster was strongly transcribed under fermentative conditions lacking external electron acceptors, reaching up to 0.2−1.9% of all transcripts, and this expression pattern closely followed that of *ldh*. In *B. subtilis* and related Bacillota, LDH expression is controlled by the redox-sensing repressor Rex, which responses to elevated NADH/NAD^+^ ratios (33–37). Notably, the G301 genome contains a Rex-binding motifs upstream of both *ldh* and the Ni-CODH/ECH operon (Fig. 6), identical to the Rex consensus sequence (36), suggesting that Ni-CODH/ECH expression in this strain is regulated by intracellular redox status. A Rex-binding motif is also present upstream of the Ni-CODH/ECH genes in *Calderihabitans maritimus* (41), suggesting that this redox-dependent regulation is potentially conserved among other hydrogenogenic CO oxidizers. During fermentation, NADH accumulation typically forces cells to balance ATP generation via acetate production with NAD^+^ regeneration via fermentation enzymes such as LDH, lowering energy yield. CO-dependent proton respiration can supply an auxiliary energy under these conditions. The activation pattern of Ni-CODH/ECH during late fermentation resembles that of FHL, which also contains a group 4a [NiFe]-hydrogenase but in association with a formate dehydrogenase that reoxidizes pyruvate-derived formate into H_2_ and CO_2_ (11, 42). As ECH complexes in Thermoanaerobacterales are likewise proposed to be Rex-regulated (10), these group 4 [NiFe]-hydrogenases may perform analogous roles in energy-conservation or mitigating acidification during reductive stress. Furthermore, as a single enzyme complex, Ni-CODH/ECH enables proton-motive force generation independently of redox carriers (e.g. quinones, NAD) and solely produces diffusible gaseous end-products (Fig. 1a). These features make its function highly resilient to variations in cellular redox state and metabolite levels.

Another key question is the source of CO for these bacteria. In natural settings, such as volcanic soils or sun-exposed/tropical soils, temperatures can reach thermophilic ranges (60 °C at the soil surface) (43), and a global geochemical study suggested that thermal degradation of organic matter is the main abiotic source of CO (44). In addition, intact soils, sediments, peatland and compost have been demonstrated to produce trace amount of CO (39, 45–48). Consistent with these environmental observations, trace accumulation of CO during anaerobic incubation of rich medium at 65 °C (Fig. S5), indicating that thermal or chemical decomposition of medium components generates CO. Notably, *P. thermoglucosidasius* Δ*cooCSF* accumulated significantly more CO than the wild-type strain, implying that the wild-type strain re-oxidized the CO produced endogenously. These findings indicate that Ni-CODH in *P. thermoglucosidasius* functions predominantly in CO consumption, contrary to the hypothesis that CO production occurs via reverse CODH activity (39). Therefore, thermophiles in soils or sediments may utilize CO from a combination of abiotic processes and metabolic byproducts. Indeed, anaerobic incubation of soils and sediments with CO yields H_2_, implying the activity of indigenous hydrogenogenic CO oxidizers (49, 50). Furthermore, hydrogenogenic CO oxidizers have been isolated or detected in environmental samples, such as marine sediments (51–53) and freshwater lake sediments (24, 54), highlighting the potential utilization of Ni-CODH/ECH in these environments. Hydrogenogenic CO oxidizers may scavenge even trace amounts of CO, aiding their survial under otherwise energy-starved conditions.

Altogether, this study elucidates a novel physiological role of Ni-CODH/ECH in enabling energy conservation and survival during electron acceptor limitation, distinct from the previously proposed CO-responsive energy conservation and/or detoxification model (3, 4, 17). Our findings demonstrate that Ni-CODH/ECH can be activated under late fermentation and is likely regulated in response to the intracellular redox state, providing new insights into how thermophilic facultative anaerobes adapt to energy scarcity. While we focused on Anoxybacillaceae, the presence of Ni-CODH/ECH in an extraordinarily phylogenetically and physiologically diverse range of bacteria and archaea suggests that this dependable form of energy conservation may supplement other microbes, as discussed previously (4, 15). Similarly to recent paradigms established for aerobic trace gas oxidation (7), we predict that CO-dependent proton respiration may be useful both as a lifeline for facultatively and obligately anaerobic bacteria experiencing electron acceptor limitation, as well as a chemolithoautotrophic or mixotrophic growth strategy for certain acetogens and methanogens with already highly constrained physiologies. Future studies should investigate the role of this complex in diverse other microorganisms and environments.

## Material and Methods

### Organisms

*Parageobacillus thermoglucosidasius* NBRC 107763^T^ (DSM 2542^T^) (29) and *Parageobacillus toebii* NBRC 107807 (55) were obtained from the Biological Resource Center of the National Institute of Technology and Evaluation (NBRC, Japan). A Δ*codh* mutant strain of *P. thermoglucosidasius* was derived from the wild-type strain and maintained in the laboratory as previously described (18). *Parageobacillus* sp. G301 (25) and *Thermolongibacillus altinsuensis* B1-1 (24) were previously isolated and preserved in our laboratory.

### Genomic analysis

A total of 378 genomes classified as Anoxybacillaceae (Taxonomy ID: 3120669) were downloaded from the NCBI reference sequence (RefSeq) database. GTDB taxonomic classifications were assigned using GTDB-Tk v2.4.0 (release R226) (56), and genome accessions with their GTDB taxonomies are listed in Table S2. The presence of CODH and hydrogenase genes was determined using DIAMOND v2.1.12 (57) with an e-value cut-off of 1 × 10^−200^. The following protein sequences were used: clade F Ni-CODH (*Parageobacillus* sp. G301, WP_285753291.1; *Rhodospirillum rubrum*, WP_200293483.1), clade E Ni-CODH (*Thermococcus onnurineus*; WP_012571978.1), and group 4a– c hydrogenase large subunit (*Parageobacillus* sp. G301 HyfG, WP_285753305.1; *T. onnurineus* MbhL, WP_012572531.1; *R. rubrum* CooH, WP_011388069.1). The phylogenetic classifications of these enzymes were based on previous studies (4, 58–60). The detection rates of CODH genes were calculated for each genus or species. A maximum-likelihood phylogenetic tree was constructed based on 120 concatenated single-copy core genes, using the GTDB backbone tree (bac120_r226). The major clades collapsed and were labeled according to the GTDB classification. Bootstrap support values (out of 100) are shown at key nodes. The phylogenetic tree was visualized using MEGA7 (61). The protein structures were predicted using AlphaFold 3 (62) and analyzed using ChimeraX (63).

### Culture experiments

Culture experiments were performed using a minimal medium as previously described (18, 27), containing per liter: 0.3 g KCl, 0.5 g NH_4_Cl, 0.1 g KH_2_PO_4_, 0.2 g MgCl_2_·6H_2_O, 0.1 g CaCl_2_·2H_2_O, 0.03 g sodium silicate, 0.1 g NaHCO_3_, 0.5 mL trace element solution SL6 (64), and 1 mL vitamin solution (65). The yeast extract concentration was set to 0.01%.

The strains were grown at 65 °C and 100 rpm in glass bottles containing 100 mL of medium, sealed with bromobutyl rubber stoppers and phenol resin screw caps. The headspace was filled with 200 mL of a gas mixture composed of 25% CO and 75% N_2_ (Kindgas, Kyoto, Japan). Sodium pyruvate (15 mM) served as the electron donor and carbon source. When specified, 25 mM MOPS-NaOH buffer (pH 6.8, 65 °C) was added to control pH changes resulting from the fermentation products.

For the transcriptomic analysis, *Parageobacillus* sp. G301 cells were cultured in a minimal medium containing 0.1% yeast extract. Either 25 mM sodium pyruvate or 20% CO was used as the electron and/or carbon source, and 0.5% sodium nitrate or 80% air (O_2_) was added as indicated. The cultures were incubated at 65 °C with shaking at 120 rpm. For nitrate-supplemented cultures, nitrate and nitrite concentrations in the liquid phase were determined using the Griess reaction with the NO_2_/NO_3_ Assay Kit CII (Dojindo Laboratories, Kumamoto, Japan).

Cell growth was monitored by measuring optical density at 600 nm (OD_600_) using an Ultrospec 2100 Pro spectrophotometer (Biochrom, Berlin, Germany). The headspace gas composition was analyzed using a GC-2014 gas chromatograph (Shimadzu, Kyoto, Japan) equipped with a thermal conductivity detector and a Shincarbon ST packed column (2.0 m × 3.0 mm; Shinwa Chemical Industries, Kyoto, Japan) using argon (Kindgas) as the carrier gas. The column temperature was programmed to increase from 40 to 200 °C. The concentrations of CO, H_2_, and CO_2_ were quantified using standard curves prepared from gas mixtures containing 0.5, 1, 5, 25, 50, and 100% CO or H_2_, and 1, 5, 10, and 20% CO_2_ or O_2_ (all > 99.99% purity; Kindgas). All standard measurements were performed in triplicates. Dissolved gas concentrations were calculated based on Henry’s law and the van’t Hoff equation (65 °C, 1 atm), using Henry’s constants of 1.4 × 10^−5^ for CO, 9.2 × 10^−6^ for H_2_, 6.9 × 10^−4^ for CO_2_, and 2.1 × 10^−5^ for O_2_ (66). Statistical analyses were performed using a linear mixed-effects model for repeated measures implemented in R v4.3.3 with the lme4, lmerTest, and emmeans packages.

### Transcriptomic analysis across 12 different conditions

Cells were harvested under 12 conditions: two carbon sources (25 mM sodium pyruvate or CO), three electron acceptors (O_2_, NO_3_^−^, and no exogenous acceptors (i.e. protons)), and two growth phases (logarithmic and stationary). Log-phase samples were collected when the OD_600_ reached 0.1, and stationary phase samples were taken at the first time point when no OD_600_ increase was observed. Cultures were centrifuged at 7,000 × g for 5 min at 25 °C, and the supernatant was discarded. Cell pellets were resuspended in 4 mL of Milli-Q water and 8 mL of RNAprotect Bacteria Reagent (Qiagen, Hilden, Germany) to stabilize the RNA and then incubated at room temperature for 5 min. Suspensions containing approximately 10^8^ cells were transferred to microcentrifuge tubes, centrifuged at 15,000 × g for 2 min at 25 °C, and stored at –80 °C until RNA extraction. Total RNA was extracted using the TRIzol Max Bacterial RNA Isolation Kit (Thermo Fisher Scientific, Waltham, MA, USA) in combination with Phasemaker Tubes (Invitrogen, Carlsbad, CA, USA), following the manufacturer’s instructions. Briefly, cell pellets were resuspended in 20 μL lysozyme solution (20 mg/mL) and incubated at 37 °C for 15 min. After incubation, 200 μL of Max Bacterial Enhancement Reagent (pre-warmed to 95 °C) was added, and the samples were vortexed vigorously to lyse the cells. TRIzol Reagent (1 mL) was then added, vortexed thoroughly, and transferred to a phasemaker tube for phase separation. DNA was removed by DNase treatment using TURBO DNase (Thermo Fisher Scientific), according to the manufacturer’s protocol. Purified RNA was further purified using the AGENCOURT RNACLEAN XP kit (Beckman Coulter, Brea, CA, USA). RNA quality was assessed using an Agilent High-Sensitivity RNA kit on an Agilent 2100 Bioanalyzer (Agilent Technologies, Santa Clara, CA, USA).

Libraries for RNA sequencing were prepared using the Stranded Total RNA Prep Ligation with Ribo-Zero Plus kit (Illumina, San Diego, CA, USA) according to the manufacturer’s instructions. Library concentrations were determined using a Qubit Fluorometer (Thermo Fisher Scientific), and fragment size distributions were evaluated using an Agilent High-Sensitivity DNA kit on an Agilent 2100 Bioanalyzer. Libraries were diluted to 4 nM, denatured with 0.2 N NaOH, and further diluted with HT1 hybridization buffer (Illumina) to a final concentration of 12 pM. Sequencing was performed on an Illumina MiSeq platform using a MiSeq Reagent Kit v3 (600 cycles). Sequencing was conducted in duplicate for stationary-phase cells grown under oxygen or in the absence of an electron acceptor and in single replicates for the other conditions, resulting in a total of 16 sequencing runs. The RNA yield and sequencing statistics are summarized in Tables S4 and S5.

Paired-end reads were processed using fastp v0.20.1 (67) to trim adapter sequences and to remove low-quality bases. Only reads with more than 80% bases with Q scores ≥30 were retained. The reads aligned to the 5S, 16S, or 23S rRNA sequences of *Parageobacillus* sp. G301 (E-value < 0.1) werewas removed using blastn from BLAST+ v2.10.1. Read quality was assessed using FastQC (https://www.bioinformatics.babraham.ac.uk/projects/fastqc/) and summarized using MultiQC v1.9 (68). Read mapping and transcript quantification were performed using Salmon v1.10.1 (69) with the coding genes of strain G301 (GCF_030268025.1) as the reference. Normalization and differential expression analysis was conducted using the edgeR package v3.30.0 (70) in R. Lowly expressed genes were filtered with filterByExpr function, and library sizes were normalized using the trimmed mean of M-values (TMM) method in edgeR. Differentially expressed genes (DEGs) were identified using quasi-likelihood F-tests implemented in glmQLFit and glmQLFTest, with a false discovery rate (FDR) < 0.01 and |log_2_ fold change (log FC)| >1. Heatmaps were generated using the R package gplots (71) using log_2_(TPM +1) values. Hierarchical clustering of log_2_-transformed TPM values was performed on genes using the Pearson correlation distance and Ward’s linkage method. Differentially expressed genes were visualized using volcano plots with logFC on the x-axis and log_10_ FDR on the y-axis. Genes of interest (e.g., *cooS* and *coxL*) were annotated manually. Plots were generated using the ggplot2 package in R (72).

## Data availability

The raw reads for RNA-seq were deposited in the NCBI/ENA/DDJB Sequence Read Archive under BioProject accession number PRJDB37689.

## Acknowledgements

Computation time was provided by the SuperComputer System, Institute for Chemical Research, Kyoto University.

## Author contributions

Y.K., M.I., S.O., and T.Y. designed research; Y.K. and S.O. performed research; Y.K. and S.O. analyzed data; Y.K., M.I., C.G. wrote the paper. The study design, experimental work, and data analysis were conducted at Kyoto University under the supervision of T.Y., with part of the final analyses performed at Monash University.

