## Supplementary Information for "Carbon monoxide-driven proton respiration enables facultative anaerobes to survive electron acceptor limitation"

### Supplementary methods and materials

#### Global detection of CO dehydrogenases and associated [NiFe] hydrogenases

The phylogenetic distribution of CODHs and group 4 [NiFe] hydrogenases was analyzed across 306,260 representative bacterial and archaeal genomes from the GlobDB r226 database (1). The archaeal and bacterial backbone trees were obtained from the GTDB r226 release. Protein sequence data were downloaded as species-level representative FASTA files and concatenated into a single searchable database. Putative CooS were identified using the Greening laboratory in-house database and DIAMOND blastp (v2.1.8) (2). All hits were filtered to retain only those with either query or subject coverage ≥80% and alignment length ≥75 amino acids. Additional filtering was applied by functional category using minimum amino acid identity thresholds of 50%. To identify potential Ni-CODH/ECH complexes, genomes were additionally screened for Group 4 [NiFe] hydrogenase genes using sequences in HydDB (3). A genome was classified as encoding a putative CODH/ECH complex when at least one [NiFe] Group 4 hydrogenase gene occurred within 15 genes upstream or downstream of a CooS homolog on the same contig. Distances between CooS and [NiFe] hydrogenase loci were computed from locus tags, and gene pairs were enumerated per genome. The resulting dataset was used for visualization of putative Ni-CODH/ECH genes distributions across microbial lineages. The archaeal and bacterial backbone trees from the GlobDB r226 release were visualized using R (v4.3.1) with the packages ggtree (4), treeio (5), ape (6), ggtreeExtra (7), cowplot (8), and colorspace (9). Phylum-level taxonomy was extracted from the GlobDB taxonomy file and visualized as colored outer rings.

#### Trace CO detection assays

To assess trace level of CO, five *Anoxybacillaceae* strains (*P. toebii*, *Parageobacillus* sp. G301, *T. altinsuensis*, *P. thermoglucosidasius* wild-type and Δ*codh* strains) were cultured in TGP medium (10), which contained per liter: 17 g tryptone, 3 g soy peptone, 5 g NaCl, 2.5 g $K_2HPO_4$, 4 mL glycerol, 4 g sodium pyruvate. Cultures were incubated at 65°C and 100 rpm in a $N_2$ headspace (100%). Each 300 mL serum bottle contained 50 mL of liquid medium and 250 mL of headspace. After 4 h of incubation, the CO concentrations in the headspace were measured using CO detector tubes (1LC; Gastec Co., Kanagawa, Japan). For time-course measurements, *P. thermoglucosidasius* wild-type and Δ*codh* strains, and *Parageobacillus* sp. G301 cells were cultured under the same conditions and headspace CO concentrations were measured at 0, 4, 19, and 25 h using the same CO detector tubes. As the CO detectors required opening the bottles for measurement, different bottles were used at each time point. All experiments were performed in triplicates.

### Supplementary figures

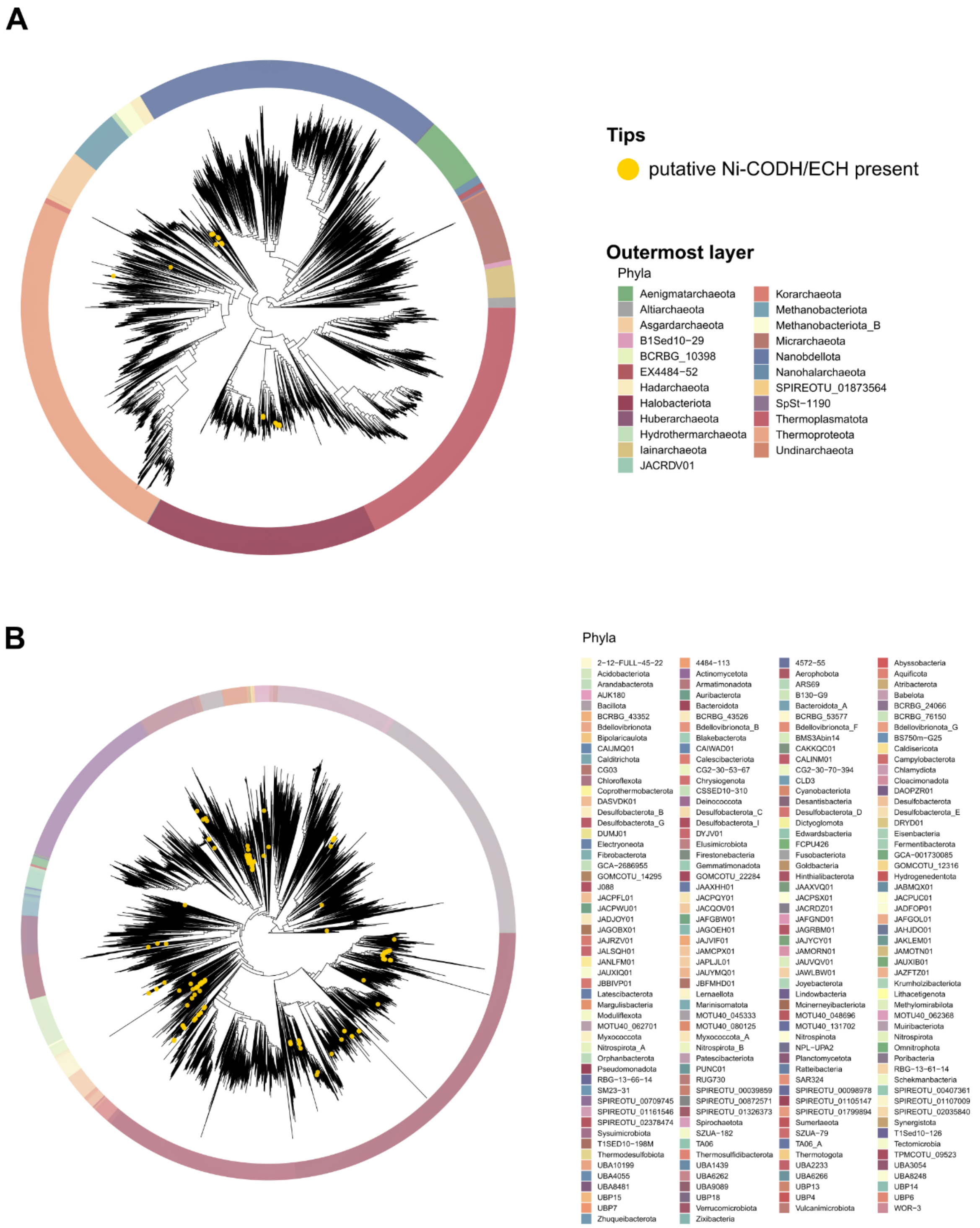


**Fig. S1. Phylogenetic distribution of putative CO dehydrogenases and putative Ni-CODH/ECH genes across 306,260 microbial genomes.**

(A) Archaeal and (B) bacterial genome trees based on the GTDB r226 backbone tree, encompassing 306,260 representative genomes from the GlobDB database (1). Outer rings indicate taxonomic affiliation at the phylum level. Yellow dots mark genomes predicted to encode a putative Ni-CODH/ECH complex, defined as those harboring a *cooS* and [NiFe] Group 4 hydrogenase gene within 15 genes.

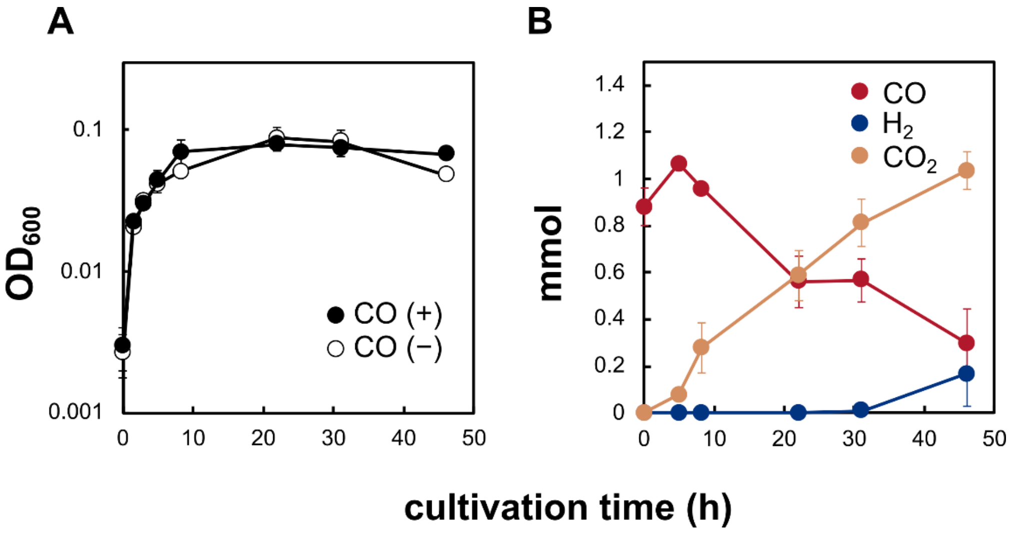


**Fig. S2. Growth and gas composition of *Parageobacillus* sp. G301 in the presence or absence of CO.**
Cultures were grown in basal medium supplemented with 5 mM sodium pyruvate under 15% $O_2$ in the presence of CO (closed symbols) or in its absence (open symbols). (A) Growth curves ($OD_{600}$). (B) Total gas amounts during the same incubations, with CO (red), $CO_2$ (yellow), and $H_2$ (blue) are shown. Data represents means of three independent biological replicates, and error bars indicate the standard error of the mean.

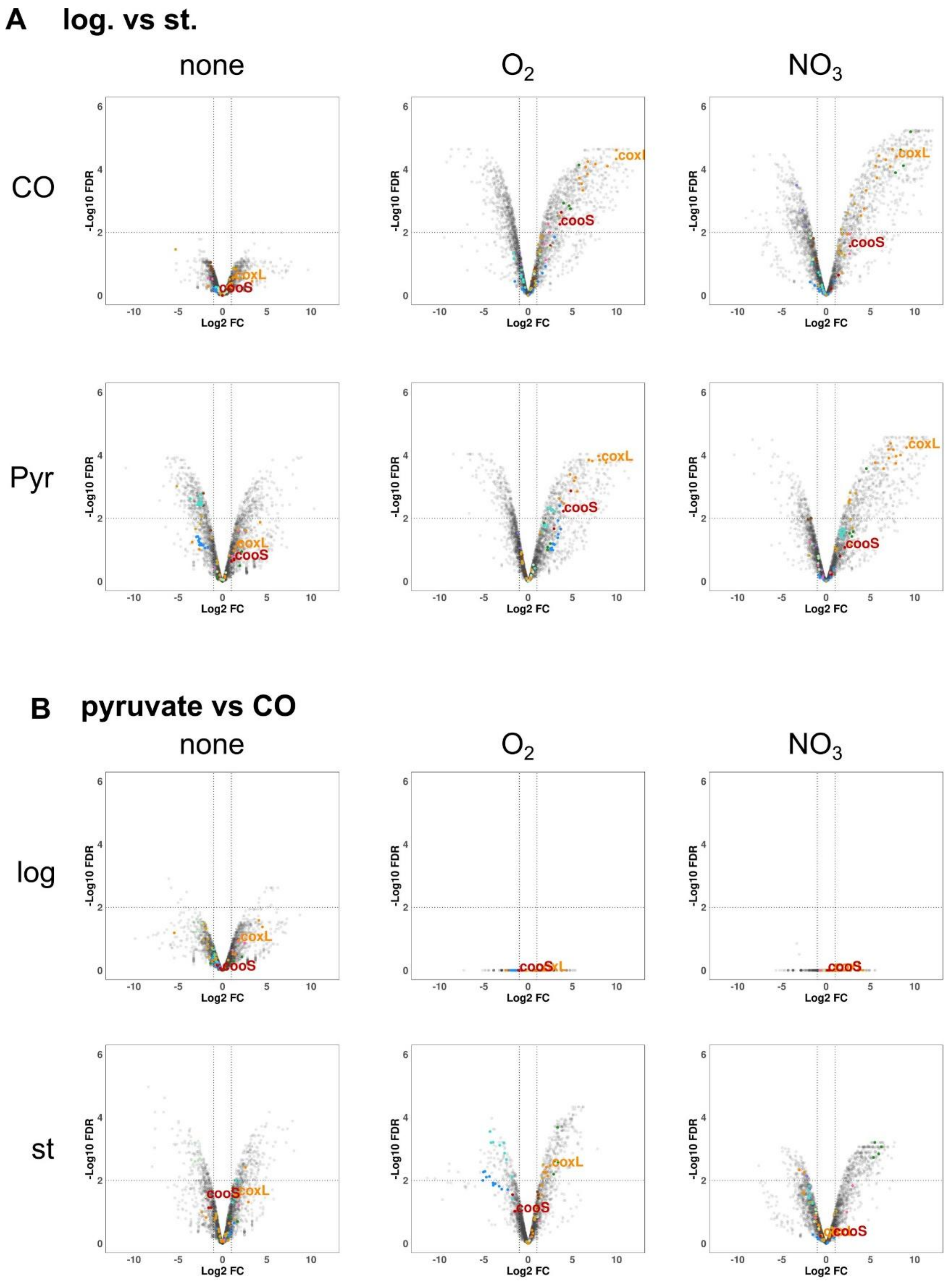


(legend on the next page)

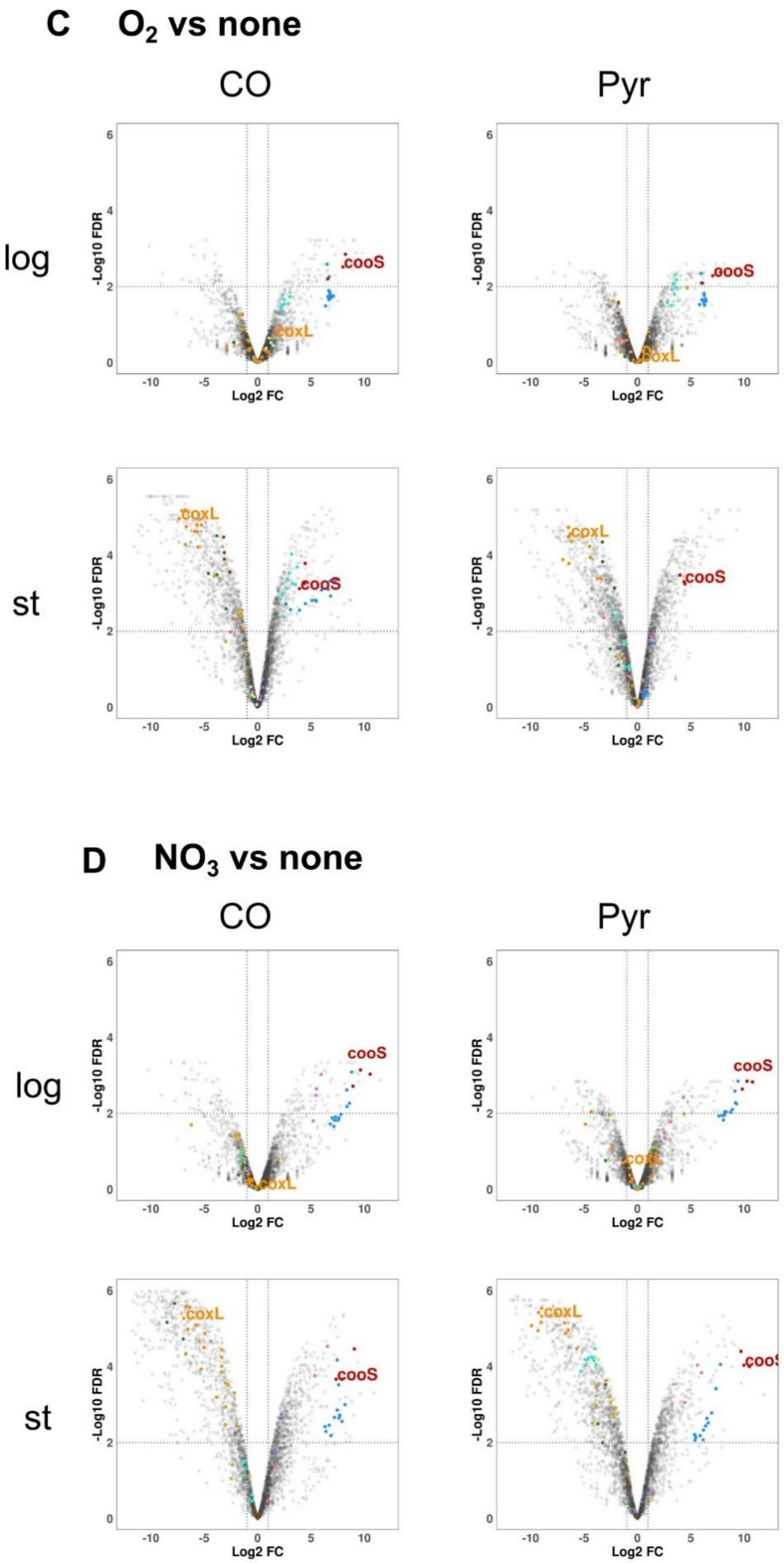


(legend on the next page)

**Fig. S3. Differential expression of *Parageobacillus* sp. G301 across substrates, electron acceptors, and growth phases.** Volcano plots show $\log_2$ fold change (x-axis) versus $\log_{10}$(FDR) (y-axis). Dotted vertical lines denote $|\log_2 FC| = 1$ and the horizontal line denotes FDR = 0.01 ($-\log_{10}FDR = 2$). Selected markers are colored by functional group (see below), and *cooS* (Ni-CODH catalytic subunit) and *coxL* (Mo-CODH catalytic subunit) are labeled when present. Gray points are all other genes. (A) Comparisons of log-phase to stationary-phase transcriptomes. (B) Comparisons of the available substrate, pyruvate or CO. (C) Electron-acceptor effect of $O_2$ relative to no addition. (D) Electron-acceptor effect of $NO_3^-$. Colors are as follows (gene groups): Ni-CODH genes (*cooCSF*), red; energy-converting hydrogenase ECH genes (*hyfB–I*, *hycH/I*, *hypA/B*), blue; Mo-CODH genes (*moc/cox*, *ctaG*), orange; cytochrome bd oxidase (*cydABCD*), pink; nitrate reductase 1 (*narGHIJ*-1), dark green; nitrate reductase 2 (*narGHIJ*-2), light green; [NiFe]-hydrogenase (*hypA–F*, *hupF/hypC*, *hycI*, *hyaABC*), turquoise; Complex I (*nuoA–N*, *ndhF*), light brown; Complex II (*sdhABC*), purple; Complex IV (*ctaCDEF*), brown.

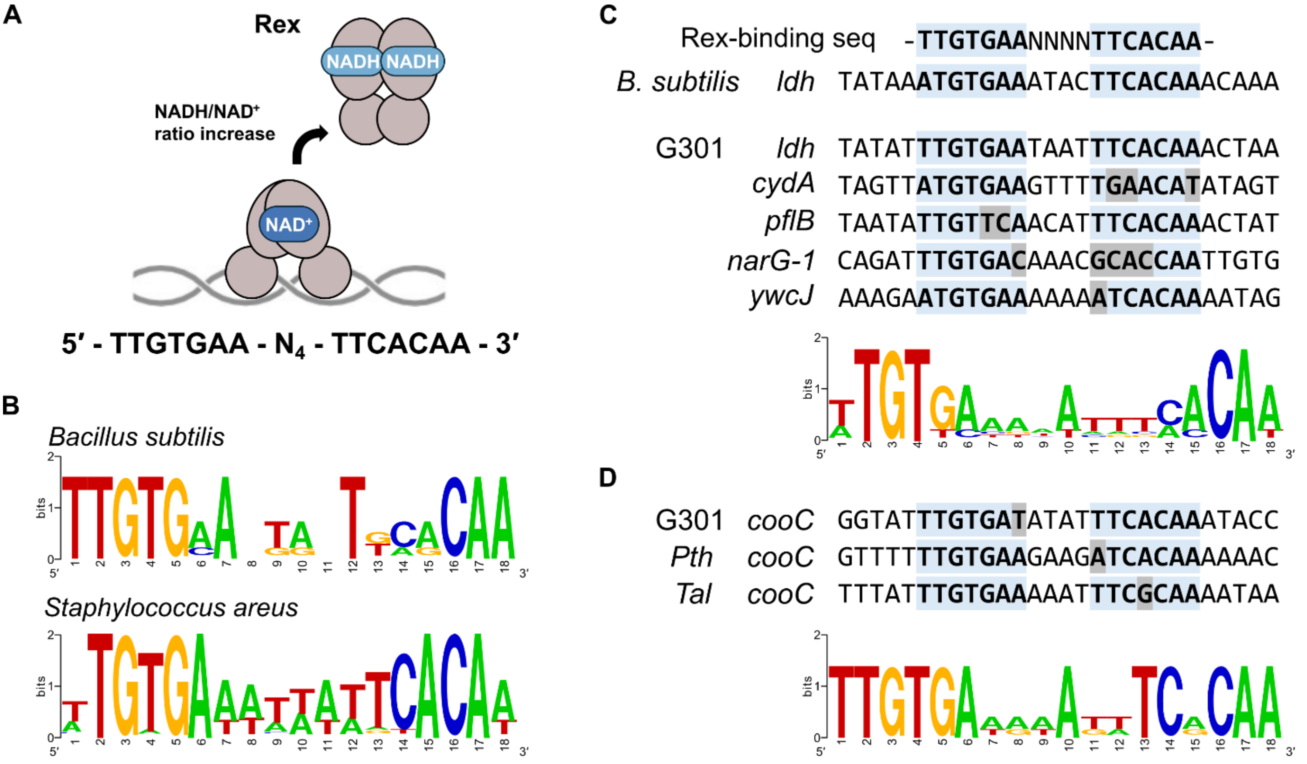


**Fig. S4 Predicted Rex-binding motifs upstream of redox-regulated genes in *Parageobacillus* sp. G301 and related species.**

(A) Schematic model of the redox-sensing transcriptional repressor Rex. Rex binds DNA as a homodimer when the intracellular NADH/$NAD^+$ ratio is low and dissociates upon NADH binding, thereby derepressing target genes (11). The consensus Rex-binding sequence (5′-TTGTGAA-$N_4$-TTCA CAA-3′) is shown below (12). (B) Experimentally validated Rex-binding motifs in *Bacillus subtilis* (13) and *Staphylococcus aureus* (14). Sequence logos were generated using WebLogo (https://weblogo.berkeley.edu/). (C) Alignment of putative Rex-binding sites upstream of potential redox-regulated genes (*ldh*, *cydA*, *pflB*, *narG*-1, and *ywcJ*) in *Parageobacillus* sp. G301. (D) Predicted Rex-binding motifs upstream of the Ni-CODH/ECH gene (*cooC*) in *Parageobacillus* sp. G301, *P. thermoglucosidasius* (*pth*), and *T. altinsuensis* B1-1 (*tal*). The sequence logo indicates the conserved Rex-binding patterns. Nucleotides matching the consensus sequence are highlighted in light blue, and mismatches are shown in gray.

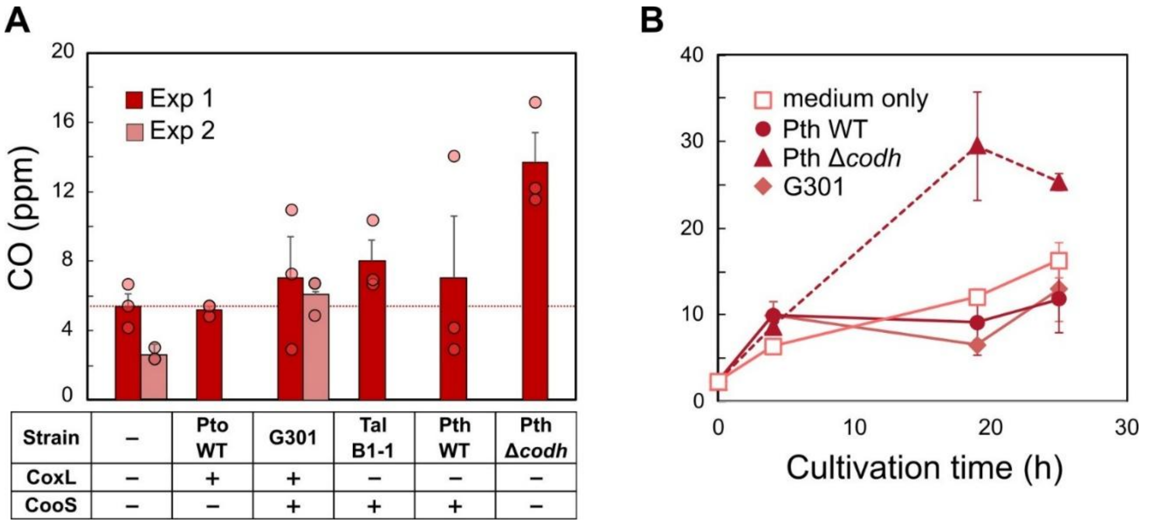


**Fig. S5 CO production by Anoxybacillaceae strains under anaerobic conditions.**
(A) The cells were cultured in TGP medium and CO concentrations measured at 4 h after inoculation. Two independent experiments (Exp 1 and Exp 2) are shown in dark and light red, respectively. Error bars represent standard deviations of biological replicates (n = 3). The dotted horizontal line indicates the CO level in the medium-only control. A table below the panel summarizes the presence (+) or absence (–) of Mo-CODH (CoxL) and Ni-CODH (CooS) in each strain. (B) Time course of CO accumulation in cultures of selected strains. CO concentrations were measured at 0, 4, 19, and 25 h. Note that each time point represents a different culture bottle. Strain abbreviations: Pto WT, *Parageobacillus toebii* NBRC 107807 (wild type); G301, *Parageobacillus* sp. G301; Tal B1-1, *Thermolongibacillus altinsuensis* B1-1; Pth WT, *Parageobacillus thermoglucosidasius* NBRC 107763 (wild type); Pth Δ*cooCSF*, CODH knockout mutant of *P. thermoglucosidasius* (Adachi et al., 2020). CO concentrations were measured using the CO detector tubes for these experiments (1LC; Gastec Co., Kanagawa, Japan). The error bars represent the standard error of the mean.
